# Pluripotency Conversion of Gene Studied from Quantum Folding Theory

**DOI:** 10.1101/018580

**Authors:** Liaofu Luo

## Abstract

The chemically and physically induced pluripotency in a stem cell is studied from the point of quantum transition between differentiation and pluripotency states in genes. The quantum folding theory of a macromolecule is briefly reviewed. The relation of protein folding rate with the number N of torsion modes participating in the quantum transition (coherence degree) is discussed and a simple statistical relationship is obtained. As a unifying approach to conformational quantum transition the folding rate formula can be generalized from protein to nucleic acid. It is found that the formula of folding rate versus N well fits RNA folding data. Then the quantum folding theory is applied to study the pluripotency conversion in stem cell. We proved that the acquisition of pluripotency is essentially a quantum-stochastic event of small-probability through the comparison of the rate with protein folding. Due to the large coherence degree of DNA and the uphill nature of torsion transition the pluripotency-acquisition rate should be small. By establishing the reaction network of the conformational change of the pluripotency genes the characteristic time of the chemical reprogramming is calculated and the result is consistent with experiments. The dependence of the transition rate on physical factors such as temperature, PH value and the volume and shape of the coherent domain of the gene is analyzed from the rate equation. Based on these studies an approach to the physically induced pluripotency with higher rate is proposed.

## 1 General formula on the conformational quantum transition rate

In quantum folding theory of macromolecule [1][2] the protein folding is regarded as a quantum transition between torsion states on polypeptide chain. The importance of torsion state can be looked as follows: as a multi-atom system, the conformation of a protein is fully determined by bond lengths, bond angles and torsion angles (dihedral angles), among which the torsion angles are most easily changed even at room temperature and can be assumed as the main variables of protein conformation (called as the slow variable of the system). Simultaneously, the torsion potential generally has several minima the transition between which is responsible for the conformational change. All torsion modes between contact residues are assumed to participate in the quantum transition coherently in protein folding. After conformation defined through torsion, the quantum transition between conformational states can be calculated. By the adiabatically elimination of fast variables we obtain the Hamiltonian *H’* describing the conformational transition of the macromolecule. Through tedious calculations of the matrix element of *H’* we obtain a general formula for the conformational transition rate [1][3]

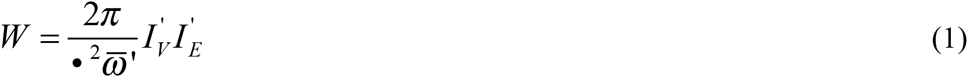

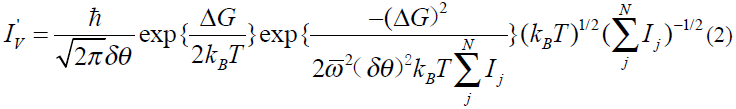

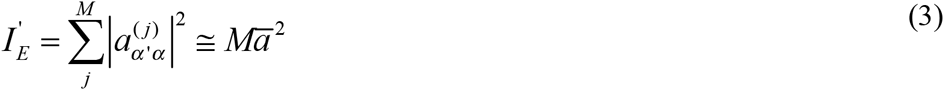

where 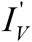 is slow-variable factor and 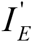 fast-variable factor, Δ*G* is the free energy decrease per molecule between initial and final states, *N* is the number of torsion modes participating in the quantum transition coherently, *I*_*j*_ denotes the inertial moment of the atomic group of the j-th torsion mode, 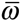 and 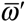 are the torsion potential parameters *ω* _*j*_ and 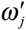 averaged over *N* torsion modes in initial and final state, respectively, *δθ* is the averaged angular shift between initial and final torsion potential, *M* is the number of torsion angles correlated to fast variables, *k*_*B*_ is Boltzmann constant and *T* is absolute temperature.

The above formula can be used for the generalized system of a macromolecule interacting with atomic groups or small molecules as long as the coordinates of atomic groups or small molecules have been added into the set of fast variables of the system [4].

## 2 Rate formula tested in protein folding

The formulae of conformational transitional rate were tested for the two-state protein folding. The statistical analyses of 65 two-state protein folding shows these formulae are in good accordance with experimental rates both in the temperature dependence (*T*-dependence) for each protein and the torsion number dependence (*N*-dependence) for different proteins [2].

Eqs(1)-(3) show that the main dependence factors of the rate is the free energy decrease (per molecule between initial and final states) Δ*G* and the fast-variable factor 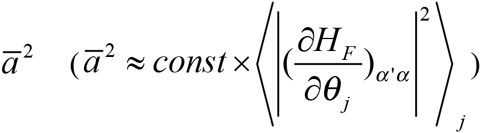, the square of the matrix element of the fast-variable Hamiltonian operator *H*_*F*_ changing with torsion angle *θ* _*j*_ averaged over *M* modes [1]. We shall study how these two factors depend on the torsion number *N*. For protein folding, RNA folding or other macromolecular conformational change not involving chemical reaction and electronic transition the fast variable includes only bond lengths and bond angles of the macromolecule. In this case a simplified expression for the fast-variable factor 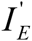 can be deduced.

Consider the globular protein folding as a typical example. Suppose the coherent transition area of the molecule is a rotational ellipsoid with semi major axis *a* and semi minor axis *b*, and the fast variable wave function *φ*_*a*_ (*x*, *θ*) a constant in the ellipsoid. The matrix element of the stretching-bending Hamiltonian is

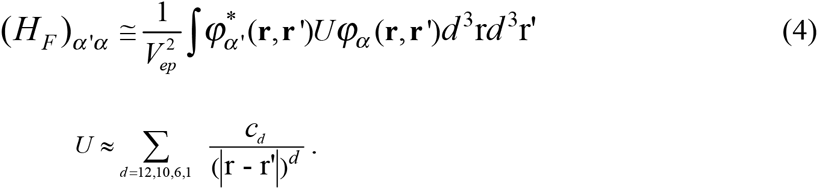

where the form of potential *U* is assumed following the general potential function for peptides [5]. It gives

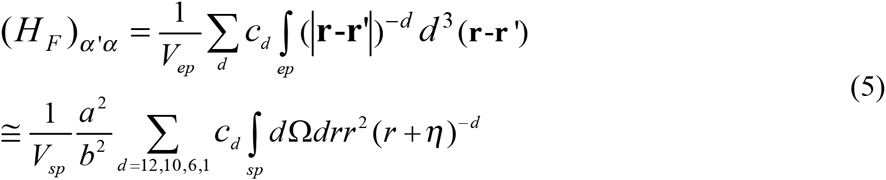

(*η* is a cutoff parameter) where 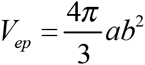 is the volume of a ellipsoid, 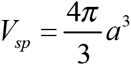 the volume of a sphere, the first integral in Eq (5) is taken over the ellipsoid and the second integral over the sphere. Because *V*_*sp*_ and *M* is proportional to *N* and the dependence of the integral 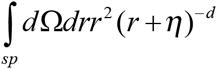 on the sphere radius is weak and can be neglected, by the consideration of the effective coupling *c*_*d*_ changing with torsion angle inversely proportional to the interacting-pair number (about *N*^−2^), from Eq (5) we estimate

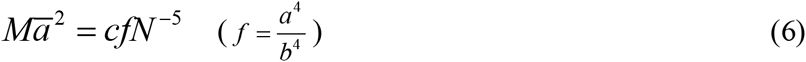

where *f* is a shape parameter and *c* an *N*-independent constant. It means the fast-variable factor 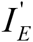 is inversely proportional to *N*^5^. This can be understood by only a small fraction of interacting-pairs (about one in *N*^2^) correlated to 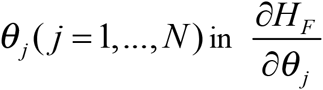. In [2] we have shown that the power law 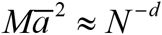 holds with *d* equal about 5.5 to 4.2 by fitting the experimental rates of 65 two-state proteins. Now the above discussion gives a theoretical demonstration of the power law. With 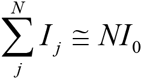 and Eq (6) inserted into Eq (1) to (3) we obtain the rate

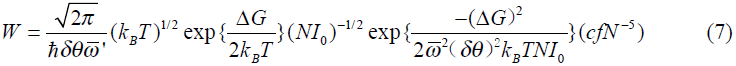

or the relation of logarithm rate with respect to *N* and Δ*G*

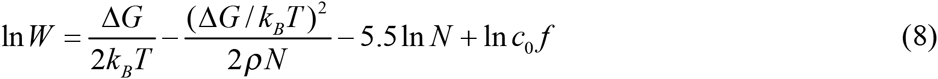

where

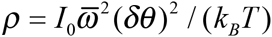 is a torsion energy-related parameter and
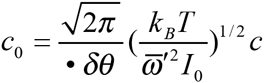 is an *N*-independent constant.

The relation of ln*W* with *N* given by Eq (8) is in good accordance with the statistical analyses of 65 two-state protein folding rates[2]. The correlation coefficient *R* between experimental logarithm folding rate and theoretical ln*W* is 0.783.

To find the relation between free energy Δ*G* and torsion number *N* we study the factor of 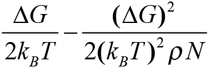 that occurs in rate equation (2) or (8). Set

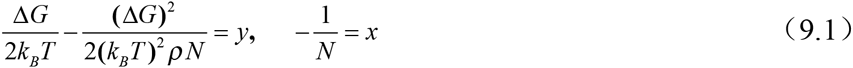

From the statistical analysis of 60 two-state proteins it gives the linear regression between y and x as shown in Fig 1,

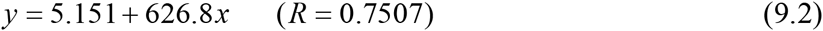

In the statistics *ρ* =0.097 as a typical value has been taken in the 60-protein set [2]. In fact, the *ρ*-values are scattered in the set. If *ρ* =0.03 is taken then the slope of the straight line in Fig 1 will decrease and Eq (9.2) is changed to

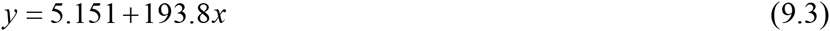

with the same correlation *R* =0.7507. However, as the variations of *ρ* are taken into account for different proteins the correlation between free energy Δ*G* and torsion number *N* can be further improved.

**Fig1.**
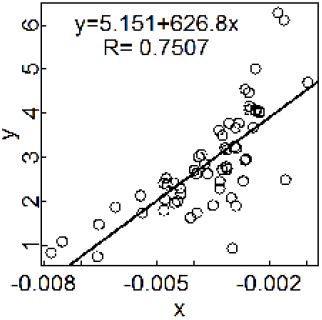
Statistical relation for 60 two-state proteins. Experimental data are taken from 65-protein set [6]. Five proteins in the set denatured by temperature have been omitted in this statistics. The figure is drawn for *π* =0.097.

About the relationship of free energy Δ*G* with torsion number *N* two statistics were done in literatures. One is based on the assumption of linear relation existing between Δ*G* and 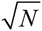 [2]. Another introduced an assumption on Δ*G vs* (*Lg* + *σ B*_*L*_ *L*^2/3^) (*L* - the length of polypeptide chain)[6]. By the statistics on two-state proteins in the same dataset we obtained the correlation between free energy and torsion-number-related quantity R=0.67 for the former and R=0.68 for the latter [2], both lower than the correlation in Fig 1. Therefore we shall use the statistical relation Eq (9) in the following studies.

In virtue of Eqs (8) (9) we obtain an approximate expression for transitional rate ln*W* versus *N* for protein folding

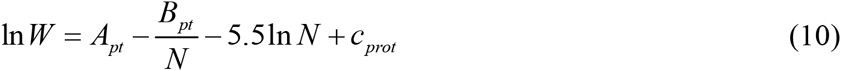

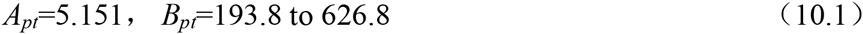

*c*_*prot*_ is an *N*-independent constant. It gives *W* increasing with *N*, attaining the maximum *W*_max_ at *N* = *B*_*pt*_ /5.5, then decreasing with power law *N*^-5.5^. The relation is consistent with the statistical analysis of the globular protein folding. In ref [2] by assuming a linear relation of free energy Δ*G* with 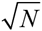, a similar decreasing relation between ln*W* and *N* is obtained but a minimum *W*_min_ is predicted therein. Since there is no evidence on the existence of the minimum, Eq (10) may be a good approximation for fitting the *N*-dependence of transitional rate.

Eqs (1-3) are introduced for calculating the conformational transition rate from quantum state *k* to *k*’. In principle, it holds equally well for the reverse process, from *k*’ to *k*. The rate of the reverse process is obtained by the replacement of Δ*G* by −Δ*G* and 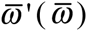 by 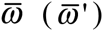 in *W*(*k →k*′). One has [1,2]

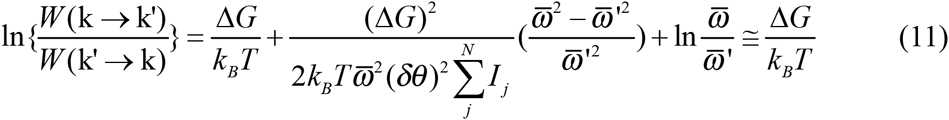

For most protein folding, Δ*G* >0. So the reverse process (protein unfolding) has a lower rate than folding by a factor exp 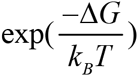. Notice that for protein unfolding Eq (10) still holds but *A*_*pt*_ should be replaced by *A*_*pt*_ = −5.151.

The quantum folding theory of protein is usable in principle for other kinds of conformational transition of macromolecule. We expect the Equations (1) to (3) are valid for nucleic acid. One may use the relation of rate *W* vs *N*, Eq(10), to analyze the RNA folding data, namely

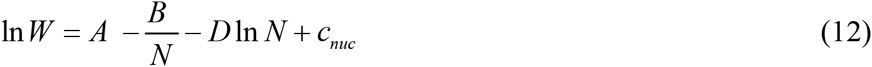

or

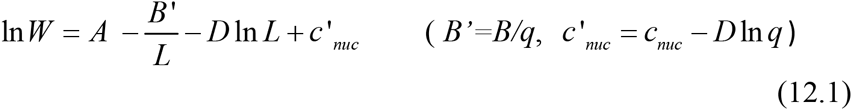

where *N=qL* (*N* proportional to *L*) is assumed, *L* is chain length of RNA and *c*_*nuc*_ - an *N*-independent constant.

In a recent literature [7] Hyeon and Thirumala (HT) indicated that the chain length determines the folding rates of RNA. They obtained a good statistical relation between folding rates *W* and chain length *L* in a dataset of 27 RNA sequences. Their best-fit result is

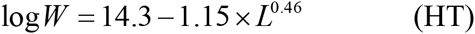

Interestingly, as a formula deduced from quantum folding theory, Eq (12) also predicts the folding rate determined by chain length. We find Eq (12.1) can fit the experimental data on RNA folding rate equally well. On the same dataset collected by [7] we obtain the best-fit *D* value is *D*_*f*_=5.619 [8], close to the theoretical prediction *D*=5.5 and the best-fit *B’* value *B*’_*f*_=61.63[8] which means *q*=3-10 if *B=B*_*pt*_ is assumed. Simultaneously, we obtain the correlation between log *W* and log *k*_*f*_ (logarithm experimental folding rate) *R*= 0.9729 by using Eq(12.1) which is same as *R*=0.9752 by using HT’s [8]. Therefore, two equations have the same overall accuracy in fitting experimental data. However, for large *L* cases the errors ∣log*W* − log *k* _*f*_ ∣calculated from Eq (12.1) are explicitly lower than those from HT. It means the folding rate lowers down with increasing *L* as *L*^−*D*^ (*D* ≅ 5.5) at large *L* (a long-tail existing in the *W-L* curve) rather than exp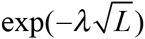 indicated by HT.

## 3 Rate formula applied to pluripotency gene

Induced pluripotent stem cells (iPSC) were firstly generated by Yamanaka in 2006. They isolated four key pluripotency genes Oct-3/4, SOX2, c-Myc and Klf4 essential for the production of pluripotent stem cells and successfully transformed human fibroblasts into pluripotent stem cells with a retroviral system [9]. The genomic integration of the transcription factors limits the utility of this approach because of the risk of mutations being inserted into the target cell’s genome. However, recently Deng et al reported in July 2013 that iPSC can be created chemically without any gene modification [10]. They used a cocktail of seven small-molecule compounds to induce the mouse somatic cells into stem cells (which they called Chemically iPSC or CiPSC) with a higher efficiency up to 0.2%. On the other hand, Su et al reported iPSC can be created directly through a physical approach [11]. They indicated that the sphere morphology helps maintaining the stemness of stem cells and proved that, due to the forced growth of cells on low attachment surface the neural progenitor cells can be generated from fibroblasts directly without introducing exogenous reprogramming factors. In the meantime, the stimulus-triggered acquisition of pluripotency was proposed tentatively and retracted soon [12,13]. More rigorous experiments with larger statistics on the physical effects on stem cells and the physically-induced pluripotency are awaited for. On the other hand, although many achievements in stem cell experiments have been made, from the point of theory, the mechanism for the chemically-physically iPSC is still one of the most puzzling and confusing problems to be understand. We shall use conformational quantum transition theory to study the pluripotency conversion and make an estimate on the probability of the conversion rate.

The fundamental processes in CiPSC include small molecules CHIR, 616452, and ESK interacting with key pluripotency-related genes Sall4 and Sox2 to enhance their expression in the early phase to activate the chemical reprogramming, and then small molecule DZNep (as a epigenetic modulator) interacting with gene Oct-4 to enhance its expression in the late phase to switch the process[10]. We suppose that when the small-molecule compounds are bound to the pluripotency gene it causes a sudden change in the molecular conformation (or shape) of the gene, namely a leap (quantum transition) from one of the torsional minima to the another of the molecule [1][4].

To apply the quantum theory of conformational change to the problem of pluripotency conversion in stem cells one should study the coherence degree *N* of a DNA molecule. For a two-state protein the torsion number *N* is calculated by the numeration of all main-chain and side-chain dihedral angles on the polypeptide chain. Typically *N* takes 100 − 300 for a typical two-state protein. For a pluripotency gene the DNA chain contains alternating links of phosphoric and sugar (deoxyribose) and every sugar is attached to a nitrogen base. Following IUB/IUPAC there are 7 torsion angles for each nucleotide, namely

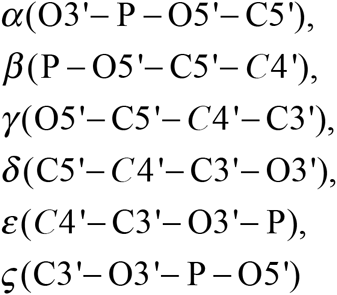

and *χ*(O4 ′– C1′– N1– C2) (for Pyrimidine) or *χ*(O4 ′– C1′– N9 − C2) (for Purine), of which many have more than one advantageous conformations (potential minima). So DNA molecule contains more torsion degrees of angles than a protein. If *W* ≈ *N* ^−5.5^ for large *N* as indicated by Eq (10) then as *N* increases tenfold, the rate *W* would decrease rapidly by a factor 3×10^5^. *N* may take a value of several thousands in the case of nucleic acid, much larger than globular protein. It affords a clue to explain why the reprogramming rate of the pluripotency-related gene is so low as compared with the protein folding rate.

We assume the differentiation state and the pluripotency state in the pluripotency-related gene is a pair of quantum states connected by torsion transition. The differentiation state has lower conformational energy (in torsion ground-state) while the pluripotency state has higher conformational energy (in torsion excited-state). The transition from pluripotency to differentiation state is a downhill reaction with free energy decrease Δ*G* >0 while the reverse transition, the acquisition of pluripotency is a process of uphill with free energy decrease Δ*G* < 0. Due to the great free energy change |Δ*G*| the difference of the rates between two process is large. This is another reason to explain why the reprogramming rate is so low. Therefore, the present theory predicts the pluripotency acquisition is a quantum-stochastic event of small-probability.

## 4 Chemically induced pluripotency calculated from quantum theory

Pluripotent stem cells were successfully induced from mouse somatic cells by small-molecule compounds. The stepwise establishment of the pluripotency circuity during chemical reprogramming was illustrated in [10]. There are seven pluripotency genes, namely Oct4, Sox2, Nanog, Sall4,Sox17, Gata4 and Gata6, involved in the pluripotency circuitry, and small molecules CHIR(C), FSK(F), 616452(6) and DZNep(Z) interacted with these genes.(Fig 2)

**Fig 2.**
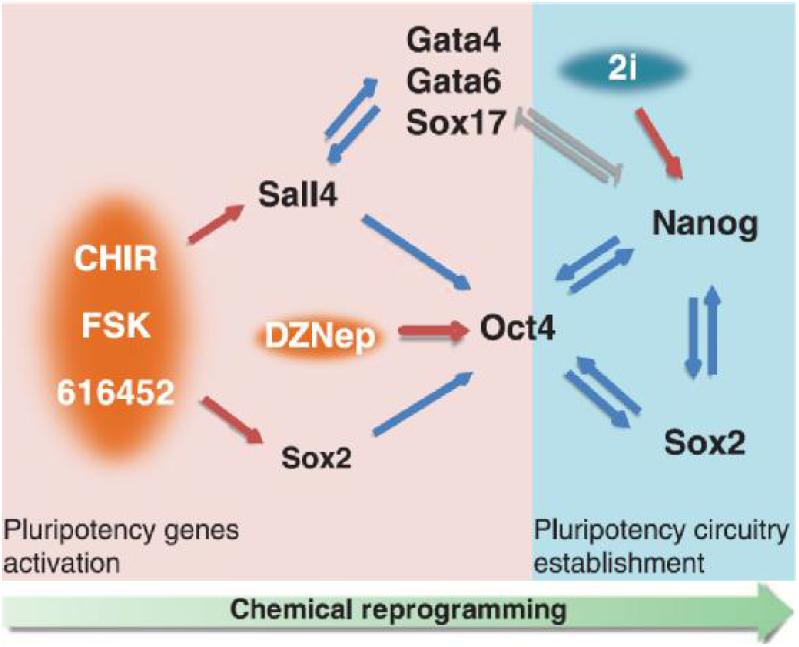
The diagram is taken from literature [10]. The schematic diagram illustrates the pluripotency circuity during chemical reprogramming. CHIR(C), FSK(F), 616452(6) and DZNep(Z) are small molecules. Oct4, Sox2, Nanog, Sall4,Sox17, Gata4 and Gata6 are seven genes involved in the pluripotency circuitry establishment.

We assume under the action of small molecules CF6 and Z the conformational transitions of pluripotency gene occur. Besides, the conformational transition also occurs in gene interaction in the circuity. Assume only the torsion transition of single gene is considered in gene interaction. That is, for a pair of interacted genes *A* and *B* one assumes the conformational transition from torsion-ground to torsion-exited state of gene A (or its reverse) in companion of gene B, namely

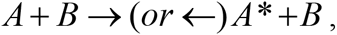

(where star * after a gene means its torsion-excited state) or the transition of gene B in companion of gene A,

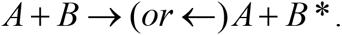

Following the pluripotency circuity illustrated in [10] we obtain a set of reaction equations of torsion transition during chemical reprogramming as follows:

1) 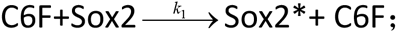
2) 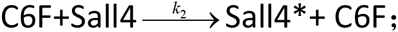
3) 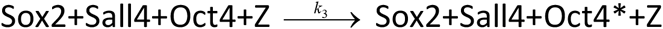 (companions Sox2 and Sall4 can be replaced by their excited states);
4) 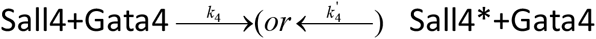 (companion Gata4 can be replaced by Gata6,Sox17 or their excited states);
5) 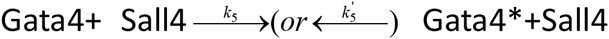 (companion Sall4 can be replaced by Nanog or their excited states);
6) 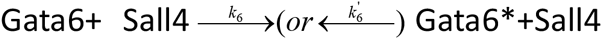 (companion Sall4 can be replaced by Nanog or their excited states);
7) 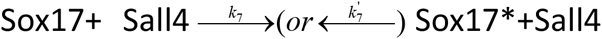 (companion Sall4 can be replaced by Nanog or their excited states);
8) 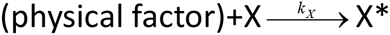(companion Gata 4 can be replaced by Gata6,Sox17 or their excited states in 2i-medium,by Oct4,Sox2 or their excited states without 2i-medium);
9) 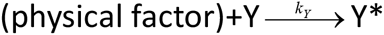(companion Nanog can be replaced by Sox2 or their excited states);
10) 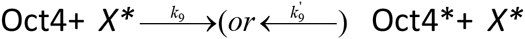(companion Nanog can be replaced by Oct4 or their excited states)

The first three equations describe the torsion transition in pluripotency genes Sox2, Sall4 and Oct4 under the action of small molecules. The next 7 equations describe the conformational transitions under gene interaction in the circuitry. The pluripotency circuitry is divided into three sub-circuits: circuit *a* of small-molecule reaction 1 to 3; circuit *b,* including gene interaction among pluripotency genes Sall4, Gata 4 (Gata6,Sox17) and Nanog described by reaction 4 to 8; and circuit *c,* including gene interaction among pluripotency genes Nanog, Oct4 and Sox 2 described by reaction 8 to 10.

From the rates {*k*_*i*_} of ten torsion transitions one can define three characteristic times corresponding to three sub-circuits,

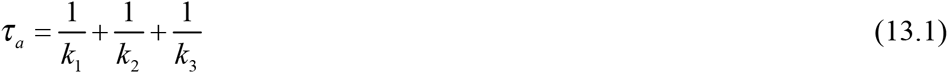

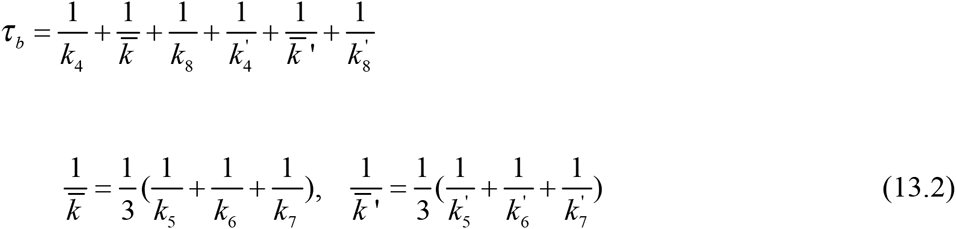

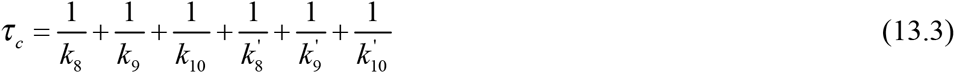

To calculate these characteristic times we estimate the rates *k*_*i*_ and *k*’_*pt*_ by using Eq (12) with *D*=5.5, *A*=-5.151 and *B*= *B*_*pt*_. By setting *k*_α_=*k*_1_,*k*_2_,*k*_3_,*k*_4_,*k*_4_^’^… … *k*_10_ or *k*_10_^’^ one calculates the ratio of gene transition rate to folding rate *k*_*pt*_ of a typical two-state protein (with torsion number *N*_*pt*_, 1/ *N* _*pt*_ = 0.003, *k*_*pt*_ = 10^4^*s*^−1^) as

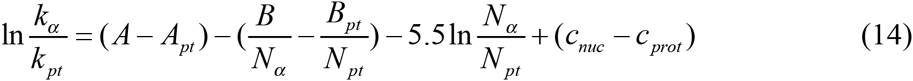

where *N*_*α*_ is the torsion number of the α-th transition. Assuming the torsion number *N*_*α*_ is proportional to the DNA sequence length of the related gene, *N*_*α*_ =*q*× (nucleotide number in α-th DNA), one has [14]

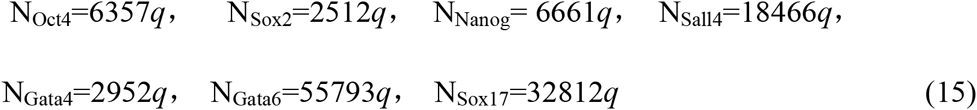

By using Eq (14) (15) we obtain

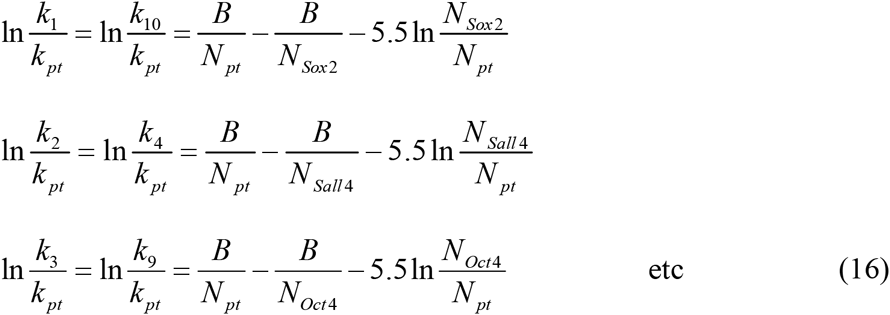

and

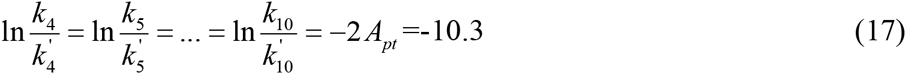

In the calculation of Eq (16) the parameters *B*=193.8 to 626.8, *c*_*nuc*_ − *c*_*prot*_ =10.3 and *q*=6 or 5 are assumed. The numerical results are given in Table 1.

**Table 1.**
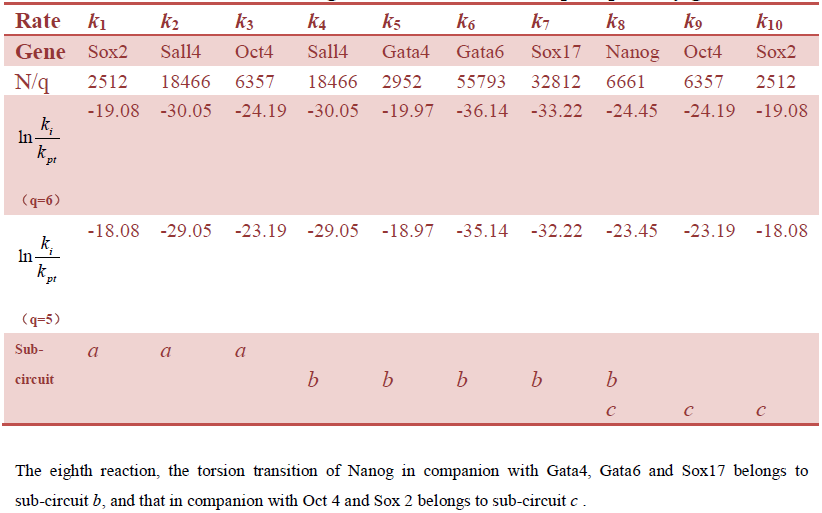
Rates of single torsion transition in pluripotency gene.

Further calculations give

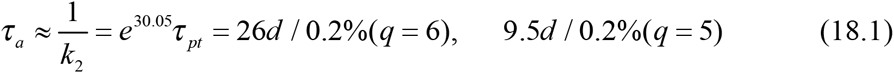

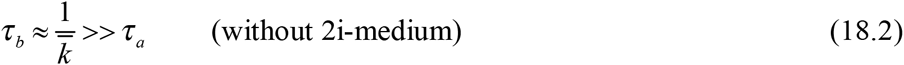

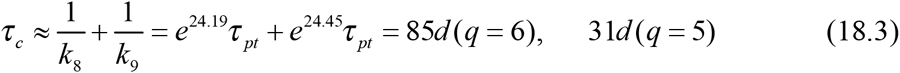

The characteristic time *τ* _*a*_ (18.1) is basically in consistency with the experimental data of pluripotent stem cells generated at a frequency up to 0.2% on day 30-40 [10] as we notice that more time is needed for post-transitional steps after the quantum transition to acquire the fully reprogrammed cell. On the other hand we notice that only after switching to 2i-medium the transitions in circuit *b* can speed up and *τ* _*b*_ < *τ*_*a*_. As *τ* _*b*_ is neglected the total pluripotency transition time *τ* equals

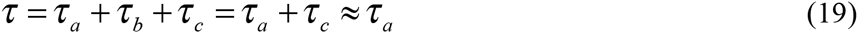

consistent with experiment.

**Note**: The above calculation shows the pluripotency transition time is in the order of several days to weeks, much longer than the protein folding time. The reason is the coherence degree *N* in DNA torsion transition much larger than that in protein folding.

The transition time *τ* _*b*_ is about 150 *τ* _*a*_ in absence of 2i medium. The time needed for full reprogramming would be too long if *τ* _*b*_ has been taken into account. However, introduce of 2i medium in circuit *b* can effectively shorten the total pluripotency transition time. Notice that the 2i medium introduced in reaction step 8 only influences the single torsion transition of Nanog gene in companion of Gata 4, Gata6 and Sox17, but has nothing to do with the same transition of Nanog in companion of Oct4 and Sox2 in circuit *c*.

In circuit *b* and circuit *c* the single torsion transition is in double directions. The reverse transition from torsion-excited to ground state is much faster (Eq 17), providing a positive feedback mechanism to the network.

Work [10] introduced small-molecule interaction to activate the pluripotency genes. Although the pluripotency transition may be speeded up by improving the small-molecule interaction, the present theory gives a lower limit for the pluripotency transition in CiPSC, namely 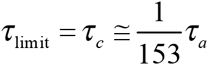.

Above points are deduced irrespective of parameter choice. The only adjustable parameters in the theory are *c*_*nuc*_ and *q* but they occur in the combination of (*c*_*nuc*_ - 5.5ln*q*). The parameter *q* means the average number of torsions in each nucleotide. The upper limit of *q* is 7 for a single chain, so the choice of *q*=6 or 5 seems reasonable because a part of torsional potentials have only one minimum. The parameter *c*_*nuc*_ is related to the free energy Δ*G* in the transition. We have assumed *c_nuc_* − *c_prot_* = 2 *A_pt_* =10.3 which is in the acceptable range of free energy change. Of course, one may use alternative parameter choices but remains *c*_*nuc*_-5.5ln*q* unaltered to obtain the same result.

We have discussed the rate of the acquisition of pluripotency from the quantum transition theory. The transition rate calculated above is for single torsion transition only. In reality the pluripotency can be acquired in multi-torsion transitions, for example, by the equation of gene interaction 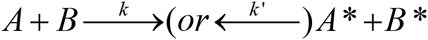. However the rate of multi-torsion transitions is much lower than the single ones and they can always be neglected.

## 5 Physically induced pluripotency inferred from the quantum theory

We shall study the physical factors influencing the rate of the acquisition of pluripotency from the quantum transition theory. The important dependence factors of the rate inferred from the theory are:

### a) Transition rate depends on temperature

Assuming the free energy decrease Δ*G* in pluripotency genes is linearly dependent of temperature *T* as in protein folding we obtain the temperature dependence of the transition rate from Eq 1 to 3,

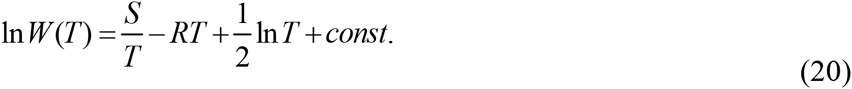

It means the non-Arrhenius behavior of the rate–temperature relationships. The relation was proved in good agreement with the experimental data of protein folding [1][2]. For a typical two-state protein the folding rate changes (increases or decreases) 2- to 7-fold in a temperature range of 40 degrees. Here we predict the same relation also holds for the reprogramming transition for the pluripotency gene..

### b) Transition rate depends on PH

Following the biochemical principle the free energy change Δ*G* of a reaction is linearly dependent on the logarithm of ion concentration [H^+^]. One has

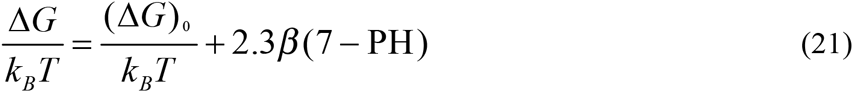

where (Δ*G*)_0_ is the standard free energy change at PH 7, *β* = O(1). The acquisition of pluripotency is an uphill process with free energy decrease Δ*G* <0. Inserting Eq (21) into Eq (1-3) we obtain

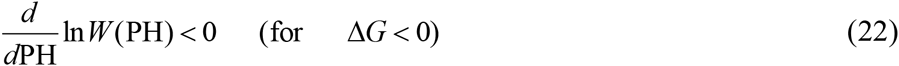

When PH decreases from 7 to 6 (or to 5, to 4, …) the logarithm rate increases by a quantity

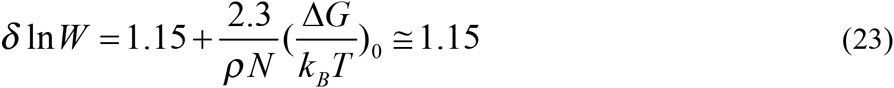

(or ≅ 2.3, 3.45, …), that means the rate increases by a factor 3.16 (or 9.97, 31.5, …). It is expected that one may more easily observe the acidity-induced pluripotency by soaking the tissues in acidic medium below PH 6.0.

### c) Transition rate depends on volume and shape of the coherent domain

As seen from Eqs (8) (12) the transition rate is strongly dependent of the coherence degree *N* of the gene. One may assume that the coherence degree is proportional to the volume *V* of the coherent domain in the gene. Therefore, from Eq (12) one has

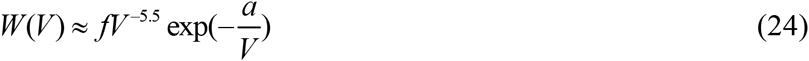

(*a*>0 is a *V*-independent constant). Eq (24) means *W* (*V*) grows as *V* decreases (in the range 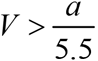). To lower down the coherence volume of the pluripotency gene a simple method is trituration of the sample under appropriate constraint.

The protein folding rate is dependent of the shape of the protein. The folding rate for a two-state protein having a quite oblong or oblate ellipsoid shape is several tenfold higher than a spheroid shape [2][6]. This can be seen by the shape parameter *f* appearing in Eqs (6) and (8). The shape parameter is another important factor to measure the coherence degree and influence the transition rate. As compared with spheroid the sphere morphology will benefit taking a larger coherence degree *N* and therefore a smaller transition rate.

Based on the above analyses we propose a physically induced pluripotency circuitry described by Fig 3. The main pluripotency circuitry is established through the cycle of Oct4-Sox2-Nanog and the latter is connected to pluripotency genes X and Y. The genes X and Y are activated through their torsion transitions under the action of physical factors.

**Fig 3.**
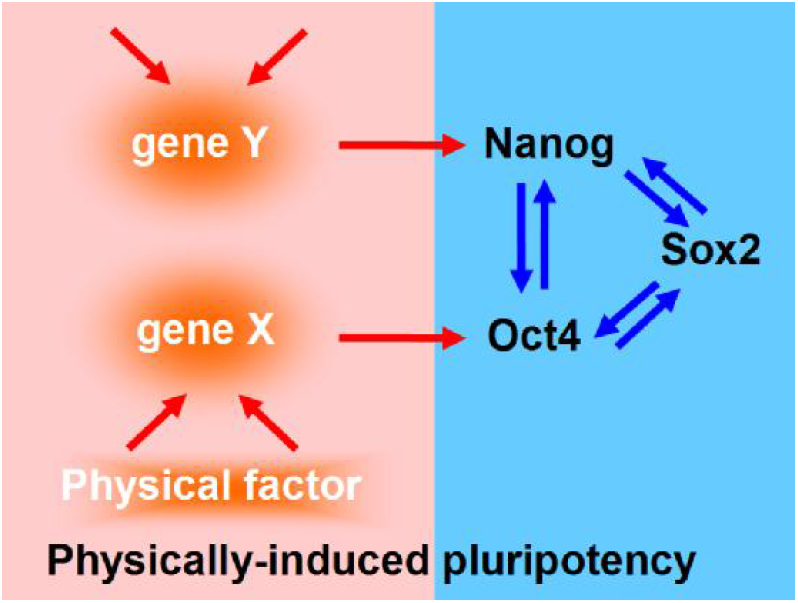
The schematic diagram illustrates the physically-induced pluripotency. The physical factor lowers down the coherence degrees in the torsion transition of gene X and Y and then establishes the pluripotency conversion in the cycle of Oct4-Sox2-Nanog.

The reaction equations are

1’) 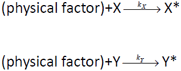
2’) 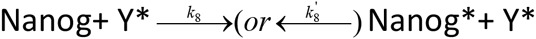(companion Y* can be replaced by Oct4 or Sox2 or their excited states)
3’) 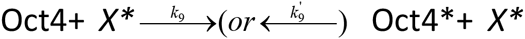 (companion *X\** can be replaced by Sox2 or Nanog or their excited states)
4’) 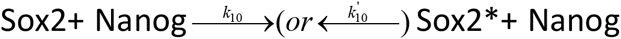 (companion Nanog can be replaced by Oct4 or their excited states)

As an example, suppose X=Sall4 and Y=Sox17. Assume the volume and shape of coherent domain of both genes are changed under physical action and as a result, both the torsion numbers of Sall4 and Sox17 decrease by a factor λ. The total pluripotency transition time of the assumed circuitry is

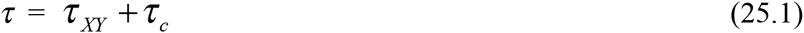

where *τ*_*c*_ is given by Eq(18.3) and

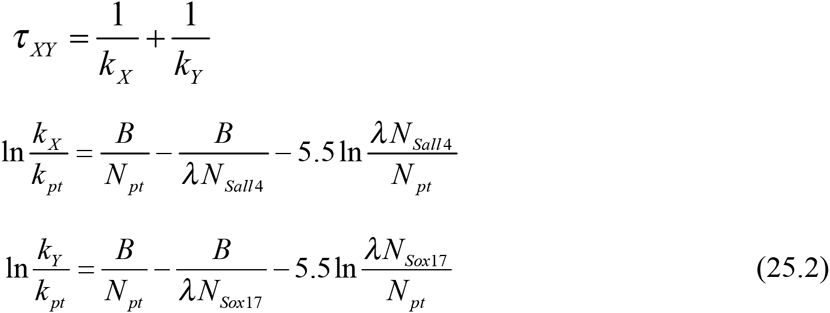

The smaller the factor λ is, the larger the rates *k*_X_ and *k*_Y_ will be. When 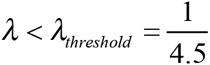 one has *τ*_*XY*_ < *τ*_*c*_ and the pluripotency transition time is dominated by *τ* _*c*_. If only X is activated by physical factor then the threshold 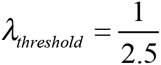. The above results are irrespective of *q* choice.

To conclude, Eqs (1) to (3) are the fundamental equations for quantum conformational transition in protein folding and pluripotency conversion of gene. For its use in pluripotency gene the key points lie in the definition of the differentiate and pluripotent state in terms of conformational state and the application of adiabatic approximation (slaving principle) to deduce the general transition formula between them. Eqs (8) (10) (12) can be used for the estimate of how the rate changes with torsion number *N* in different processes. The important thing is: In spite of the various complex pathways in the stem cell development the pluripotency genes always have the torsion quantum transition as a key step in the reprogramming of stem cells. The quantum peculiarity of the conformational change determines the pluripotency acquisition is a stochastic event of small probability. Simultaneously, the stability of quantum state gives a natural explanation on the maintenance of the stemness of stem cells.

Moreover, we have shown the physical factors such as temperature and acidity can change the rate of torsion transition. So, low PH exposure and heat shock may have observable effect in the pluripotency conversion. However, we found the most effective approach to increase the torsion transition rate is to lower down the coherence degrees of the gene. Based on the proposed model the physically induced pluripotency transition rate can, in principle, increase more than 150 times as compared with that in the known small-molecule induction approach [9].

## Acknowledgement

The author thanks Drs Jun Lu and Judong Zhao for their statistical analyses on protein and RNA folding, Drs Yulai Bao and Yennie Cao for their helps in literature and data searching.

